# High-flow Nano-chromatography Columns Facilitate Rapid and High-quality Proteome Analysis on Standard Nano-LC Hardware

**DOI:** 10.1101/501908

**Authors:** Christopher S. Hughes, Hans Adomat, Stephane LeBihan, Colin Collins, P.H. Sorensen, Gregg B. Morin

## Abstract

Optimizing the acquisition of proteomics data collected from a mass spectrometer (MS) requires careful selection of processed material quantities, liquid-chromatography (LC) setup, and data acquisition parameters. The small internal diameter (ID) columns standardly used in nano-chromatography coupled MS result in long per injection overhead times that require sacrifices in design of offline-fractionation and data acquisition schemes. As cohort sizes and the numbers of samples to be analyzed continue to increase, there is a need to investigate methods for improving the efficiency and time of an acquisition (LC + MS). In this work, the ability to improve throughput in single runs or as part of an in-depth proteome analysis of a fractionated sample using standard LC hardware is investigated. Capitalizing on the increased loading capacity of nanochromatography columns with larger IDs, substantially improved throughput with no reduction in detection sensitivity is achieved in single-injection proteome analyses. An optimized 150 μm ID column setup is paired with an offline fractionation-concatenation scheme to demonstrate the ability to perform in-depth proteome analysis on-par with current state-of-the-art studies, while minimizing sample loading overhead. Together, these data demonstrate an easy and effective means to improve sample analysis throughput with no reduction in data quality using an approach that is applicable to any standard nano-LC hardware.

## Introduction

Maintaining an optimum balance between liquid chromatography (LC) and mass spectrometer (MS) acquisition parameters is essential to maximize the content and quality of the resulting data without requiring inordinate overall analysis times. For the examination of small molecule analytes, ‘standard’ high-performance liquid chromatography (HPLC) systems coupled to MS are often used. These HPLC systems conventionally operate in analytical flow rate regimes (>100μL/min) and are paired with large bore (>1mm internal diameter, ID) chromatography columns (5-15 cm) packed with sub-2μm particles that combine to provide ultra-fast and high-resolution separation of analyte mixtures. This type of HPLC configuration provides advantages in throughput, robustness, reproducibility, and long-term stability. However, when coupled to MS, measurement sensitivity is not fully optimized for analytes with limited amounts of available material due to the large post-column elution volumes (sample dilution) and less efficient ionization. With significant optimization, these systems have been successfully applied to proteome analysis with MS ^1–4^, but the required complexity in achieving these performance levels with this type of setup limits their widespread use.

To increase measurement sensitivity for the analysis of proteomes, nano-versions of these HPLC systems are frequently employed. These nano-LC systems typically operate in low flow rate regimes (<1μL/min) and are coupled with small bore (<200μm ID), long chromatography columns (15 – 50cm) packed with sub-2μm particles. As a result of the small column bores and particle sizes, nano-LC systems are usually capable of providing reliable mobile phase flow and mixing at extremely high pressures (e.g. 1,000 bar). This type of nano-LC configuration is ideal for peptide samples as the nano-chromatography columns used feature low post-column elution volumes and greater ionization efficiency which maximizes sensitivity with electrospray MS ^5^. However, nano-columns can be difficult to prepare, expensive to purchase, can have short lifetimes, are more susceptible to peak broadening due to sample overloading, and are prone to clogging. In addition, slow equilibration and sample loading steps that stem from pressure and flow rate limitations when using small bore columns can result in substantial per-injection overhead time.

To overcome challenges associated with injection overhead, deep proteome analyses can be performed in using single injection (‘single-shot’) acquisition without offline pre-fractionation ^6^. Single-shot analyses are advantageous as there are fewer upstream processing steps and only a single nano-LC injection is required per sample, minimizing overhead time. However, for complex samples, single-shot analyses require high efficiency nanochromatography columns that can be challenging to prepare at long lengths and can exhibit substantial backpressure during operation. Furthermore, restrictions in loading capacity can limit the dynamic range of MS detection in single-shot runs. As an alternative, offline fractionation can be used to circumvent these issues by separating a complex peptide mixture into multiple lower complexity samples that are then individually analyzed by the MS ^7^. Aside from requiring more input material and processing steps, the primary pitfall of offline fractionation is the increased sample loading overhead associated with multiple nano-LC injections. As a result of this overhead, offline fractionation users will frequently separate samples initially to a larger set that are concatenated down to a smaller final injection group (e.g. 96 fractions concatenated to 12) ^8^. The concatenation scheme used needs to obtain a balance that maximizes fraction ‘uniqueness’ while minimizing overhead related to the number of injections required to analyze a final set. Recently, an innovative nano-LC hardware platform (Evosep One) was introduced that circumvented injection overhead issues ^9^. In this system, each sample is preloaded onto individual C18 matrix tips that are robotically positioned in the analytical column flow path for analysis, allowing the bulk of the required loading to be performed offline. As a result of this unique operation, LC loading overhead can be reduced to less than 2 minutes per injection depending on the parameters used, eliminating one of the major hurdles facing offline fractionation workflows. However, it remains unclear whether the underlying principles of this system can be adapted to improve the performance of standard nano-LC hardware.

In this work, the performance of standard nano-LC hardware (Easy nanoLC 1100 – 1200 series) is evaluated when using nano-chromatography columns packed into capillaries with a range of internal diameters (ID). Although other studies have compared nano- and micro-flow column performance ^1,4^, there is currently limited data on the use of larger nano-flow columns (100 μm – 200 μm ID) with integrated electrospray tips on current generation LC hardware across a range of injection quantities and applied to the in-depth analysis of a proteome. The experiments described here illustrate the ability to substantially reduce nano-LC injection overhead on standard hardware without sacrificing measurement sensitivity. The ability to obtain proteome coverage to a depth of >140,000 unique peptides (>9,700 proteins) using a 150 μm ID column with ion trap and Orbitrap MS acquisition coupled to an offline fractionation workflow is demonstrated. Importantly, these data illustrate the benefit of achieving optimum MS operation efficiency by adjusting the fraction-concatenation scheme to maximize fraction diversity and the nano-LC column configuration to minimize overhead. These performance gains can be achieved on any nano-LC system, without the requirement for bespoke hardware or other custom software control packages.

## Experimental Section

### Cell culture and harvest

HEK293T cells were grown in DMEM supplemented with 10% fetal bovine serum (FBS) in a 37°C and 5% CO_2_ incubated environment. Cells were harvested at 80% confluency using trypsin. A total of 1×10^8^ cells were grown, harvested, and stored at −80°C as a single cell pellet until use.

### Detergent-based protein isolation, reduction, and alkylation

The harvested and frozen cell pellet was reconstituted in 1mL of a gentle lysis buffer containing: 50 mM HEPES pH 7.3 (Fisher Scientific, CAT#BP299-1), 60 mM KCl (Sigma, CAT#P9541), 5 mM MgCl_2_ (Sigma, CAT#M8266), 0.1% (v/v) NP40 (Sigma, CAT#NP40), and 0.5X EDTA-free complete protease inhibitor cocktail (Sigma, CAT#04693132001)). To the lysis mixture, 20 μL of Benzonase (Sigma, CAT#E8263) was added and the tube was incubated at 37°C for 15 minutes. After incubation, 1 mL of secondary lysis solution (4% (wt/vol) SDS (Fisher Scientific, CAT#BP1311), 10 mM dithiothreitol (Bio-Rad, CAT#1610611)) was added and the mixture incubated at 60°C in a ThermoMixer (Eppendorf) for 30 minutes at 1,000rpm. Iodoacetamide (Bio-Rad, CAT#1632109) was added to a final concentration of 20 mM and the mixture incubated for 30 minutes at 24°C in the dark. Additional dithiothreitol was then added to a final concentration of 20mM to quench the reaction. Protein concentration was measured using a BCA Assay (Thermo Fisher Scientific, CAT#23225). The prepared mixture was stored at −80°C until use.

### SP3 protein clean-up and tryptic digestion

For processing to peptides, six separate aliquots of 100 μg aliquots of measured protein were prepared in 100 μL volumes. SP3 processing was carried out as described previously ^10–12^. Briefly, a 1:1 combination of two different types of carboxylate-functionalized beads (Sera-Mag Speed Beads, GE Life Sciences, CAT#45152105050350 and CAT#65152105050350), were twice rinsed in water using a magnetic rack prior to processing. Beads were added to protein mixtures to achieve an estimated concentration ratio of 1:10 (μg of protein: μg of SP3 beads). To initiate binding, ethanol was added to achieve a specific final concentration (50% by volume). Tubes were incubated at room temperature for 5 minutes with mixing at 1,000 rpm in a ThermoMixer. Tubes were placed in a magnetic rack and incubated for 2 minutes at RT. The supernatant was discarded and the beads rinsed 3 times with 180 μL of 80% ethanol by removing the tubes from the magnetic rack and resuspending the beads by pipette mixing. For elution, tubes were removed from the magnetic rack, and beads resuspended in 200 μL of 50 mM HEPES, pH 7.3 containing an appropriate amount of trypsin/rLysC mix (1:50 enzyme to protein concentration) (Promega, CAT#V5071). Tubes were sonicated briefly (30 seconds) in a bath sonicator and incubated for 14 hours at 37°C in a ThermoMixer with mixing at 1000 rpm. For peptide recovery, the tubes were briefly (30 seconds) sonicated in a bath sonicator and subsequently centrifuged at 20,000 g for 2 minutes. Tubes were then placed on a magnetic rack and the supernatant recovered for further processing.

### High-pH reversed phase fractionation

High-pH reversed phase analysis was performed on an Agilent 1100 HPLC system equipped with a diode array detector (254, 260, and 280 nm). Fractionation was performed on a Kinetix EVO-C18 column (4.6 mm x 250 mm, 2.6 μm core shell, Phenomenex, CAT#00G-4725-E0) fitted with a KrudKatcher Ultra precolumn (Phenomenex, CAT#AF0-8497). Columns were heated to 50°C using Hot Sleeve™ column ovens (Analytical Sales and Services). Elution was performed at a flow rate of 1mL per minute using a gradient of mobile phase A (20mM ammonium bicarbonate, pH 8) and B (acetonitrile). The gradient was 5% B for 5 minutes, to 8% B in 2 minutes, to 16% B in 13 minutes, to 40% B in 25 minutes, to 80% B in 1 minute, held at 80% for 4 minutes, to 5% B in 1 minute, and a final reconditioning at 5% for 9 minutes (60 minutes total runtime). From 5 – 45 minutes, fractions were collected every 25 seconds (96 total fractions) and concatenated into 24 final samples (fraction 1 = A1, C1, E1, G1, fraction 2 = A2, C2, E2, G2, etc…). Fractions were dried in a SpeedVac centrifuge and reconstituted in 0.1% trifluoroacetic acid prior to further clean-up.

### Peptide clean-up prior to MS

Peptides were desalted and concentrated by C18 solid phase extraction (SPE) using 96-well Slit Plates (Glygen). For Slit Plate clean-up, wells in 10 μL plates (Glygen, CAT#S2C18) were rinsed twice with 200 μL of acetonitrile (Sigma, HPLC-grade, CAT#34998) with 0.1% TFA by centrifuging for 30 seconds at 200 g. Wells were then rinsed twice with 200 μL of water (Sigma, HPLC-grade, CAT#34877) with 0.1% TFA by centrifuging for 60 seconds at 200 g prior to sample loading. Samples were loaded by spinning for 60 seconds at 200 g. Loaded samples were rinsed three times with 0.1% formic acid (200 μL per rinse) and eluted directly into 96-well autosampler plates with 200 μL of 60% acetonitrile containing 0.1% formic acid. Eluted peptides were concentrated in a SpeedVac centrifuge (Thermo Scientific) and subsequently reconstituted in 1% formic acid (Thermo Scientific, LC-MS grade, CAT#85178) with 1% dimethylsulfoxide (Sigma, CAT#D4540) in water.

### Mass spectrometry data acquisition

Analysis of peptides was carried out on an Orbitrap Fusion Lumos MS platform (Thermo Scientific). Samples were introduced using an Easy-nLC 1200 system (Thermo Scientific). The Easy-nLC 1200 system was plumbed with a single-column setup using a liquid-junction for spray voltage application. The factory 20 μm ID x 50 cm S-valve column-out line was replaced with a 50 μm ID x 20 cm line to reduce backpressure during operation at high flow rates. Columns were packed in 50 μm, 100 μm, 150 μm, and 200 μm ID capillaries using a nano-LC pump to push the beads from a reservoir. Briefly, fritted 200 μm ID reservoir capillaries are filled with beads using a pressure bomb (nanoBaume, Western Fluids). Packed capillaries are back-flushed using an Eksigent NanoLC pump into the final desired column capillaries at a flow rate of 1 μL/min (60% acetonitrile, 40% water), and the pressure ramped to >600 bar to compress the material. All columns were prepared with fritted nanospray tips (formamide and Kasil 1640 in a 1:3 ratio) using a laser puller instrument (Sutter Instruments). All columns were packed to a length of 20 cm with 1.9 μm Reprosil-Pur C18 beads (Dr. Maisch) in an acetone slurry.

The analytical columns were connected to the Orbitrap Fusion Lumos MS using a modified version of the University of Washington Proteomics Resource (UWPR) Nanospray ion source (http://proteomicsresource.washington.edu/protocols05/nsisource.php) combined with column heating to 60°C using a 15 cm AgileSLEEVE column oven (Analytical Sales & Service). Prior to each sample injection, the analytical column was equilibrated at 400 bar for a total volume of 3 μL. After injection, sample loading was carried out for a total volume of 6 μL at a pressure of 400 bar. The injection volumes for all samples was 2 μL across all analyses. For the column ID comparisons, 30 minute runs were performed with a gradient of mobile phase A (water and 0.1% formic acid) from 3 – 7% B (80% acetonitrile with 0.1% formic acid) over 2 minutes, to 35% B over 19 minutes, to 95% B over 0.25 minutes, hold at 95% B for 3 minutes, to 3% B in 0.25 minutes, and holding at 3% for 5.5 minutes. Flow rates were adjusted based on the column ID: 50 μm = 200 nL/min, 100 μm = 400 nL/min, 150 μm = 800 nL/min, and 200 μm = 1500 nL/min. For the proteome analyses, 60 minute runs were performed with a gradient of mobile phase A (water and 0.1% formic acid) from 3 – 7% B (80% acetonitrile with 0.1% formic acid) over 2 minutes, to 35% B over 41 minutes, to 95% B over 0.25 minutes, hold at 95% B for 3 minutes, to 3% B in 0.25 minutes, and holding at 3% for 5.5 minutes with a flow rate of 800 nL/min.

Data acquisition on the Orbitrap Fusion Lumos (control software version 3.1.2412.17) was carried out using a data-dependent method with MS2 in the Orbitrap. The Fusion Lumos was operated with a positive ion spray voltage of 2400 and a transfer tube temperature of 325°C. The default charge state was set as 2. Survey scans (MS1) were acquired in the Orbitrap at a resolution of 60K, across a mass range of 400 – 1200 m/z, with an S-Lens RF lens setting of 60, an AGC target of 4e5, a max injection time of 54ms in profile mode. For dependent scans (MS2), monoisotopic precursor selection, charge state filtering of 2 – 4 charges, and dynamic exclusion for 15 seconds with 10ppm low and high tolerances were used. A 1 m/z window was used prior to HCD fragmentation with a setting of 35%. MS2 data acquisition carried out in the Orbitrap used a 30K resolution, a fixed first mass of 110m/z, an AGC target of 1.2e5, and a max injection time of 54ms in centroid mode. MS2 data acquisition carried out in the ion trap used a Rapid scan speed, a fixed first mass of 110m/z, an AGC target of 1e4, and a max injection time of 54ms in centroid mode. Injection of ions for all parallelizable time was turned off for both Orbitrap and ion trap MS2 acquisitions.

### Mass spectrometry data analysis

All data files were processed with RawTools (version 1.4.0) to generate instrument operation reports, performance metrics, and MGF outputs ^13,14^. For MGF creation, mass and charge state recalibration with RawTools was enabled (-puxmR –chro 1B flags). For peptide matching as part of the quality control search functionality of RawTools, the X!Tandem (version 2017.2.1.4) search engine was used. The ‘-N’ flag of RawTools was set to 5,000 to randomly select this number of MS2 spectra from each file for searching. MS2 data were searched on-the-fly by RawTools and X!Tandem against a UniProt human proteome database (version 2018_10) containing common contaminants (21,098 total target sequences). Decoy proteins were generated on-the-fly by RawTools for target-decoy analysis. RawTools automatically reads the instrument configuration and adjusts the mass accuracy settings based on the determined mass analyzer. Precursor accuracy settings were set at 10ppm, with product accuracy at 0.05 Da for the MS2 Orbitrap data. Carbamidomethylation of cysteine was set as a fixed modification and oxidation of methionine variable. Trypsin enzyme rules with a total of 2 missed cleavages allowable was specified. X!Tandem results were filtered by RawTools by taking the 95^th^ percentile decoy score and keeping all target hits above this value.

For searching RawTools generated MGF outputs for all acquired samples and published data (PXD006932, PXD010393) ^15,9^, a combination of SearchCLI (version 3.3.9) ^16,17^ and PeptideShakerCLI (version 1.16.29) ^18^ was used. For PXD006932 ^15^, the 46 fraction data acquired at 15,000 resolution were used. The data were pulled from this alternative repository because although used in PXD010393, this more recent repository was missing the raw data for the 33^rd^ fraction. All searches used a combination of X!Tandem (version 2015.12.15.2)^19^, MS-GF+ (version 2018.04.09) ^20^, and Tide (version 3.0.17109) ^21^ algorithms. MS2 data were searched against a UniProt human proteome database (version 2018_10) containing common contaminants (The Global Proteome Machine cRAP sequences – https://www.thegpm.org/crap/) that was appended to reversed sequences generated using the -decoy tag of FastaCLI in SearchCLI (42,196 total sequences, 21,098 target). Identification parameter files were generated using IdentificationParametersCLI in SearchCLI specifying precursor and fragment tolerances of 20ppm and 0.5 Da (ion trap MS2) or 0.05 Da (Orbitrap MS2), carbamidomethyl of cysteine as a fixed modification, and oxidation of methionine and acetylation of protein N-term as variable modifications. Trypsin enzyme rules with a total of 2 missed cleavages allowable was specified.

All SearchCLI results were processed into PSM, peptide, and protein sets using PeptideShakerCLI. Error rates were controlled in PeptideShakerCLI using the target-decoy search strategy to determine false-discovery rates (FDR). Hits from multiple search engines were unified using posterior error probabilities determined from the target-decoy search strategy. Results reports were exported from PeptideShakerCLI using the ReportCLI with numeric values provided to the -reports tag to provide the ‘Certificate of Analysis’, ‘Default Protein Report’, ‘Default Peptide Report’, and ‘Default PSM Report’. All results (PSM, Peptide, Protein) were filtered to provide a final FDR level of <1%. Final mzid files output from PeptideShakerCLI used MzidCLI with the default parameters.

### General statistical parameters

In all boxplots, center lines in plotted boxes indicate the median, upper and lower lines the 75^th^ and 25^th^ percentiles, and upper and lower whiskers 1.5X the interquartile range.

### Data and code availability

The mass spectrometry proteomics data were deposited to the ProteomeXchange Consortium (http://proteomecentral.proteomexchange.org) via the PRIDE partner repository ^22,23^ with the dataset identifier PXDXXXX (NOTE – the data have been submitted to PRIDE on December 12, 2018. They have not returned an accession yet). The repository contains all raw data, search results, and sequence database files.

R Notebook files (markdown and html format) detailing data analysis and figure creation for this manuscript are all freely available on GitHub: https://github.com/chrishuges/ColumnIDs_JPR-2019.

## Results and Discussion

Determining the optimum setup when coupling nano-LC to MS depends on the design and goals of a given experiment. One of the primary bottlenecks that results in substantial overhead in proteomics experiments is the long times required to equilibrate and load sample into small ID nano-chromatography columns due to the extreme backpressures that limit the possible flow rates for these steps. The simplest way to reduce backpressure without sacrificing separation efficiency is to increase the ID of the column. Although previous reports have demonstrated the severe sensitivity drop resulting from increasing column ID, they primarily focused on 1 mm or greater diameters. We reasoned that increasing column ID in the range of 150 μm – 200 μm using capillaries with integrated electrospray tips would provide an improved balance between loss of sensitivity and higher throughput afforded by higher flow rates.

### High-flow nano-chromatography columns provide enhanced throughput with minimal sensitivity loss

To first test the general performance of columns prepared with different IDs (each column was packed to 20 cm with 1.9 μm C18 beads), a set of triplicate injections of 250 ng of HEK293 tryptic peptide digests was analyzed with each setup. A specific flow rate was used for each column (50 μm = 200 nL/min, 100 μm = 400 nL/min, 150 μm = 800 nL/min, 200 μm = 1,500 nL/min) and the required volume for equilibration and sample loading were kept constant to facilitate accurate examination of ‘dead times’ for each injection. As the 50 μm column contains less volume compared to those with larger IDs, we aimed to minimize the equilibration volume (3 μL) to ensure the overhead analyses were not substantially biased against the smaller columns. The same gradient elution steps and MS acquisition conditions were used across all injections. The sample injection volume was kept constant across all samples (2 μL). All equilibration and loading steps were performed at a maximum pressure of 400 bar for two primary reasons: 1. The Easy nanoLC system used exhibits substantially improved reliability operating at these medium pressures; 2. Using a lower pressure limit ensures the approach detailed here can be extended to a wide range of UHPLC systems, including those that cannot reach ultra-high-pressures (1,000 bar).

Based on the calculated injection cycle dead times, the significantly improved throughput of the larger ID chromatography columns was clearly observable (Figure 1a). Injection cycle overhead time was calculated as the amount of time required for column equilibration, sample injection, and loading (from the start of the injection cycle to the first MS scan event). The 50 μm column required an average of 51 minutes to complete an injection cycle at 400 bar, a duration longer than the MS acquisition itself (mean 50 μm = 51 minutes, 100 μm = 11 minutes, 150 μm = 8 minutes, 200 μm = 8 minutes). This is largely due to the low flow rate resulting from the high backpressure generated by this column (200 nL/min @ 400 bar). The larger ID columns afforded higher flow rates at the pressure settings used (100 μm = 1.1 μL/min, 150 μm = 1.9 μL/min, 200 μm = 2.2 μL/min), enabling more rapid column equilibration and sample loading, ultimately reducing injection overhead time. However, quality examination of the data files for each injection using RawTools reproducibly showed the reduced performance for the higher ID columns in terms of precursor signal, MS2 intensity, and electrospray stability (Figure 1b-d). This in turn resulted in a reduction in the numbers of MS2 scans acquired and the rate of identification from these spectra (Figure S1a-b). Across all tested IDs, the 200 μm column generally performed substantially worse than the others tested in these experiments.

**Figure 1.**
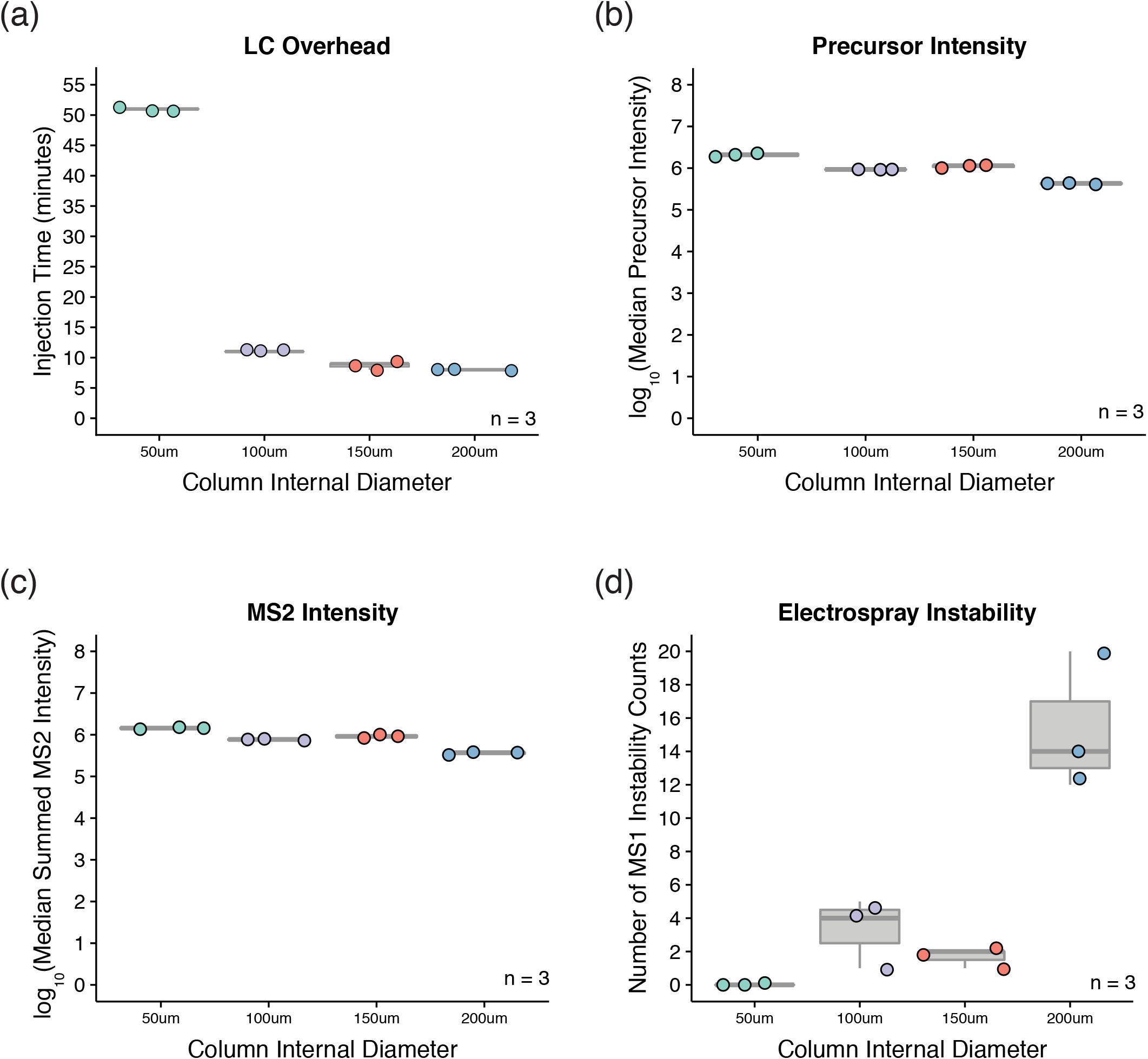
Nano-chromatography columns with larger IDs improve throughput with minimal sensitivity loss. Peptides derived from an HEK293 lysate were subjected to injection replicate (n = 3) analyses on an Easy nanoLC system equipped with columns of increasing ID (50 μm, 100 μm, 150 μm, 200 μm). Each sample was analyzed by an Orbitrap Fusion Lumos for 30 minutes using the same gradient conditions. **(a)** Boxplot depicting the time in minutes required to perform the injection, equilibration, and sample loading steps prior to MS analysis. **(b)** Boxplot of the logarithm transformed median intensity values of all MS1 precursors that triggered MS2 scans in each run. **(c)** Boxplot of the logarithm transformed median of the summed intensity values from each of the MS2 scans acquired across each analysis. **(d)** Boxplot of the numbers of electrospray instability events observed in each analysis. Electrospray instability is defined as the number of MS1 scans whose neighbor differs in signal by >10-fold.

To quantify the overall performance of the different columns, two separate metrics indicative of MS operation efficiency were calculated, injection and acquisition efficiency. Injection efficiency was calculated as the proportion of time spent performing MS acquisition relative to the total time for an analysis (Injection efficiency = (MS acquisition time / (MS acquisition time + LC overhead time)) * 100). The injection efficiency provides insight into how much time per injection was being spent performing actual data acquisition. The other metric, acquisition efficiency, was calculated as the proportion of MS2 scans acquired relative to all collected scan events (Acquisition efficiency = (Number of MS2 scans / Total Number of Scans) * 100). Acquisition efficiency provides insight into how much time during the MS analysis was actually spent performing dependent analysis on parent ions of interest. As a final output, the product of the injection and acquisition values (injection efficiency * acquisition efficiency) / 100) designated MS Efficiency (MSE) was then used to quantify overall performance. From these calculations, the substantial drop in MSE was further observed for the 50 μm column (Figure 2a-c). The observed drop was mainly driven by the injection efficiency, where the large overhead time contributed to a significant reduction in this value. Taking these trends into account, the 150 μm ID column was selected for further analysis as it offered the best balance of overhead and MSE, while not suffering substantially from negative aspects like poor electrospray stability.

**Figure 2.**
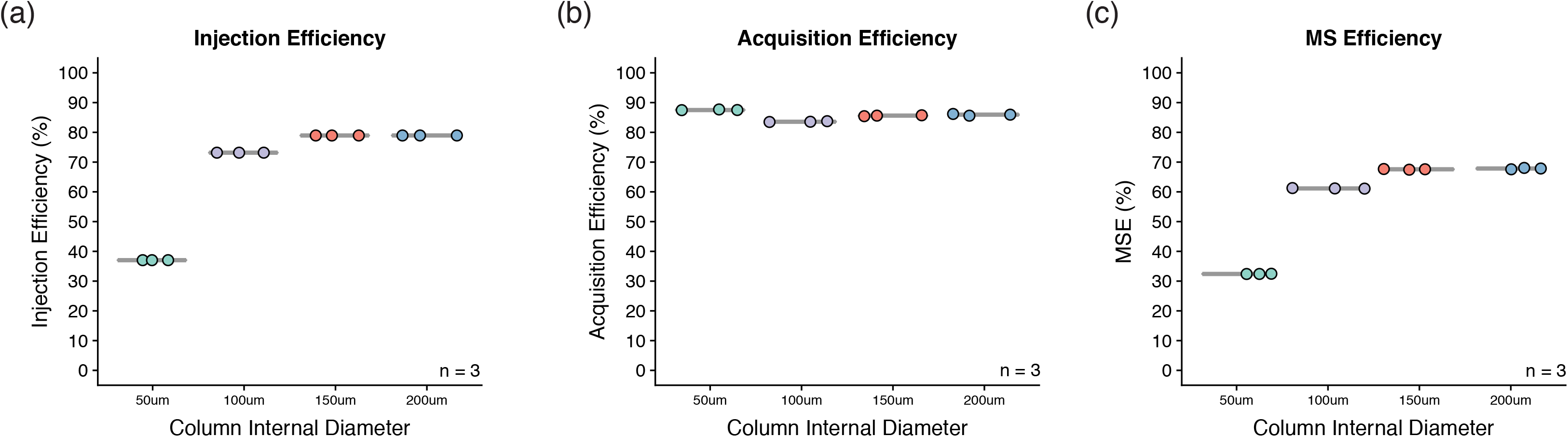
The efficiency of MS acquisition is substantially improved for analyses undertaken with larger column IDs. Measurements of MS efficiency were calculated based on the injection replicate (n = 3) analyses of an HEK293 peptide sample on an Easy nanoLC system equipped with columns of increasing ID (50 μm, 100 μm, 150 μm, 200 μm). **(a)** Boxplot depicting the injection efficiency across the different column IDs. Injection efficiency is defined as the amount of time required for MS analysis as a proportion of the total time required for sample acquisition (LC overhead + MS analysis time). **(b)** Boxplot depicting the acquisition efficiency across the different column IDs. Acquisition efficiency is defined as the proportion of all scans that were acquired that were dependent (MS2) events. **(c)** Boxplot depicting the MSE values across the different column IDs. MSE is defined as the product of the injection and acquisition efficiency scores divided by 100.

A noteworthy advantage of a large column ID is the ability to increase loaded sample quantities without negatively impacting chromatography performance. To test the potential for recovery of sensitivity on the 150 μm compared to the 50 μm column, various injection amounts were analyzed in triplicate (injection amounts = 250 ng, 500 ng, 1000 ng, 2000 ng, injection volume maintained at 2 μL for all samples). Examination of this data revealed the precursor signal, MS2 intensities, and MS2 identification rate levels could be restored to be on par with, or exceeding, those for the 50 μm column (Figure 3a-c). Importantly, these improvements did not come at the cost of chromatography performance, as there was only a minimal rise in observed peak width as injection amount was increased (Figure 3d). These increases also did not come at the cost of MSE, where scores were maintained at 69% across all of the tested injection amounts. Taken together, these data indicate that increasing column ID (along with a subsequent flow rate increase) substantially improves sample analysis throughput but decreases the overall sensitivity of the MS analysis. However, the loss of sensitivity can be mitigated by increasing the load amount to where the larger ID columns meet or exceed the performance of those with small IDs.

**Figure 3.**
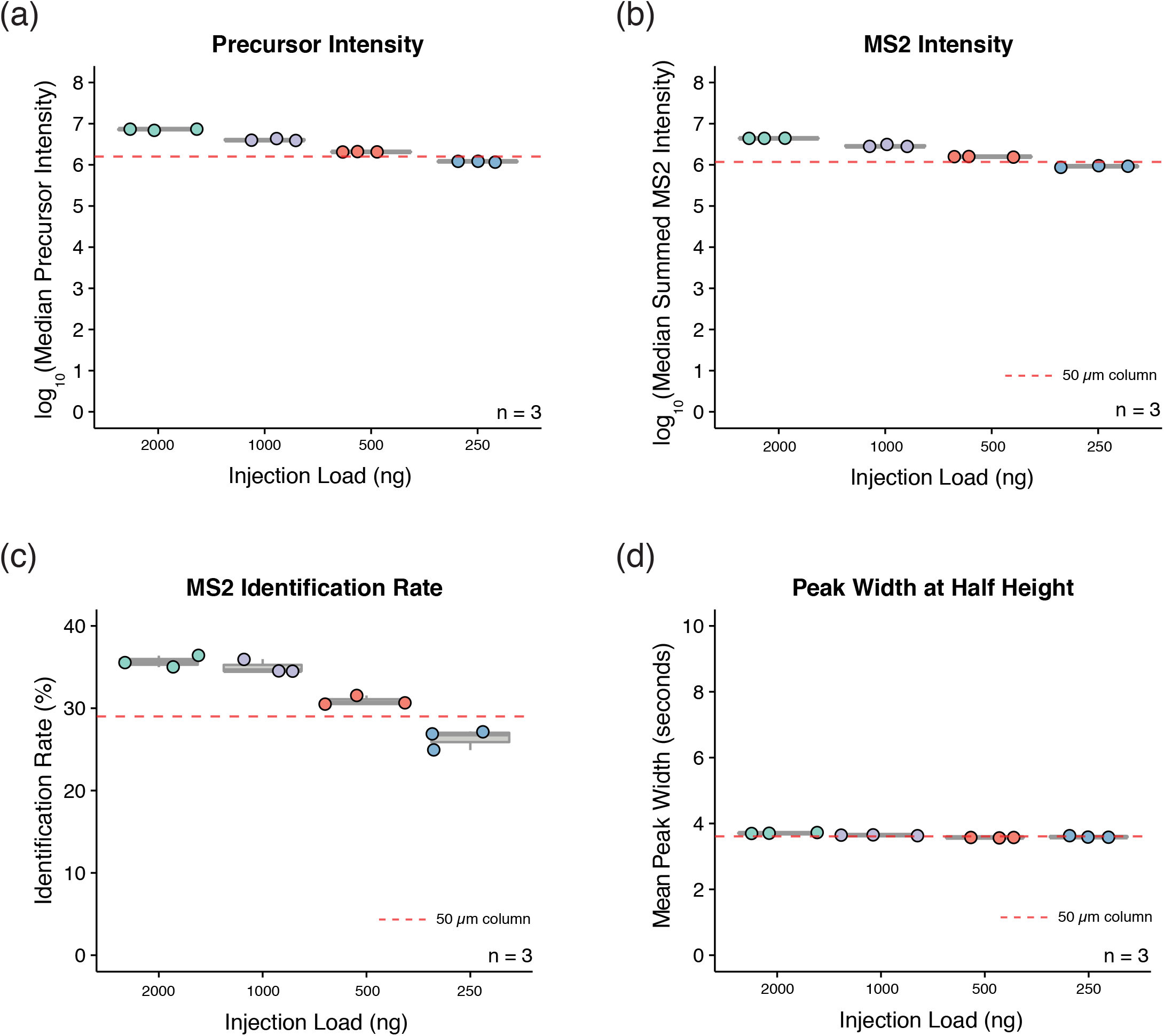
The reduced sensitivity when using larger column IDs can be recovered by injection of more peptide material. Peptides derived from an HEK293 lysate were subjected to injection replicate (n = 3) analyses on an Easy nanoLC system equipped with a 150 μm ID column. Each sample was analyzed by an Orbitrap Fusion Lumos for 30 minutes using the same gradient conditions. The injection quantity was scaled to provide on-column amounts of 250 ng, 500 ng, 1,000 ng, and 2,000 ng (all using a 2 μL injection). **(a)** Boxplot of the logarithm transformed median intensity values of all MS1 precursors that triggered MS2 scans in each run. **(b)** Boxplot of the logarithm transformed median of the summed intensity values from each of the MS2 scans acquired across each analysis. **(c)** Boxplot depicting the identification rate of peptides from MS2 spectra across the different injection amounts. Values are derived from RawTools QC analysis using X!Tandem on a random selection of 5,000 MS2 spectra per file. **(d)** Boxplot depicting the observed mean of the half-height peak widths for all precursors observed that triggered MS2 events in each analysis. In all of the above plots, the red dashed line indicates the mean of the value reported for the 50 μm ID column (250 ng load) for the same parameter.

### High-flow nano-chromatography columns facilitate rapid analysis of fractionated proteomes

In order to obtain enhanced coverage of a complex proteomics sample when using MS, offline fractionation of peptide digests is often used. To maximize the efficiency of the subsequent MS analysis, samples are typically fractionated into a larger set (e.g. 96 fractions) and then concatenated down to a smaller group (e.g. 12 concatenated fractions). This concatenation serves two purposes: 1. To ensure a uniform representation of peptides across the entire gradient is present in any given fraction (minimizing the possibility of empty elution windows); 2. To minimize the number of injections required to analyze the overall fraction set (to minimize LC overhead). An optimum balance between these two purposes is achieved when uniform elution of peptides is observed across an entire gradient for each fraction with the uniqueness in the composition maintained, while avoiding sacrificing extensive analysis time due to LC overhead.

Recently, the Evosep One system was described and applied to the analysis of offline-fractionated peptides on a Q-Exactive HF-X MS system ^9^. A peptide mixture generated from a HeLa cell lysate was separated offline into 46 fractions (no concatenation) and analyzed using either an Easy nanoLC or an Evosep One, on the same MS. The data clearly demonstrated the enhanced performance of the Evosep One system, where just 18.4 hours (16.1 hours of gradient time, 3 minutes of LC overhead per fraction) was required to analyze the entire fraction set, compared with 28.3 hours (14.6 hours of gradient time, 17.8 minutes of LC overhead per fraction) for the Easy nanoLC system. An efficiency of MS utilization metric that is the same as the ‘injection efficiency’ value calculated by the authors of the above study revealed that the Evosep system substantially outperformed the standard Easy nanoLC system (88% vs. 52% injection efficiency). However, examination of MS1 chromatograms and MS2 scan triggering revealed the presence of extended empty elution windows in the early fractions in the set for either system (Figure S2a-d), as described previously ^14^. These data suggest that by combining a fraction-concatenation scheme with a larger ID nano-LC column to increase nano-LC injection efficiency, the overall MSE could potentially be improved.

To test whether the analysis efficiency could be improved on an Easy nanoLC system using the larger 150 μm ID column setup described in this work, a peptide mixture from HEK293 cells was separated offline into 96 fractions and concatenated down to 24 final samples for MS analysis. A final set of 24 fractions was selected as a balance between reducing uniqueness due to concatenation and the associated LC overhead from the number of injections required. Each of the 24 fractions was analyzed for 60 minutes on an Orbitrap Fusion Lumos system using both the ion trap and Orbitrap for the MS2 acquisitions. Although isobaric tags were not utilized here, when using the Orbitrap for MS2 detection the resolution, fill time, and AGC parameters were set to be compatible with TMT analysis (30,000 at m/z 200, 54ms, 1.2e5) to enable extension of these results to these multiplexing experiment types. Based on the analyses above that indicated that the injection overhead for the 150 μm column was 8-minutes, the elution gradient and MS acquisition were set to 52 minutes to result in a final analysis time per fraction of 60 minutes (24 hours total acquisition time, including LC overhead).

Database search analysis of the acquired data revealed the detection of 148,294 unique peptides (9,700 proteins) from the ion trap data, and 151,388 unique peptides (9,749 proteins) from the Orbitrap. Re-analysis of the Easy nanoLC and Evosep One data described above using the same search pipeline resulted in 158,788 peptides (8,717 proteins) and 159,416 peptides (8,941 proteins) from the two systems (Figure 4a). RawTools analysis of the data revealed a mean MSE across all fractions of 71% (ion trap) and 75% (Orbitrap) for the 150 μm ID Easy nanoLC setup employed here (Figure 4a). The 150 μm ID setup outperformed the mean MSE calculated for the 75 μm ID Easy nanoLC (45%) and was comparable to the MSE for the Evosep One (75%) from the previously published data (Figure 4a). Importantly, the MSE for the 150 μm ID Easy nanoLC setup was consistent across the fractions (Figure 4b), owing largely to using the concatenation approach designed to minimize the presence of empty elution windows. Taken together, these data demonstrate the ability to achieve substantially higher analysis throughput of fractionated and concatenated material without sacrificing overall identifications through the use of a larger ID nano-LC column setup. However, it is important to mention that the Evosep system could achieve further potential gains via the use of fraction-concatenation. A theoretical calculation of a final set of 24 concatenated sample injections with an adjusted 60-minute run (57 minutes run time, 3 minutes of LC overhead based on published data) using an Evosep One system showed the injection efficiency would approach an impressive 95%.

**Figure 4.**
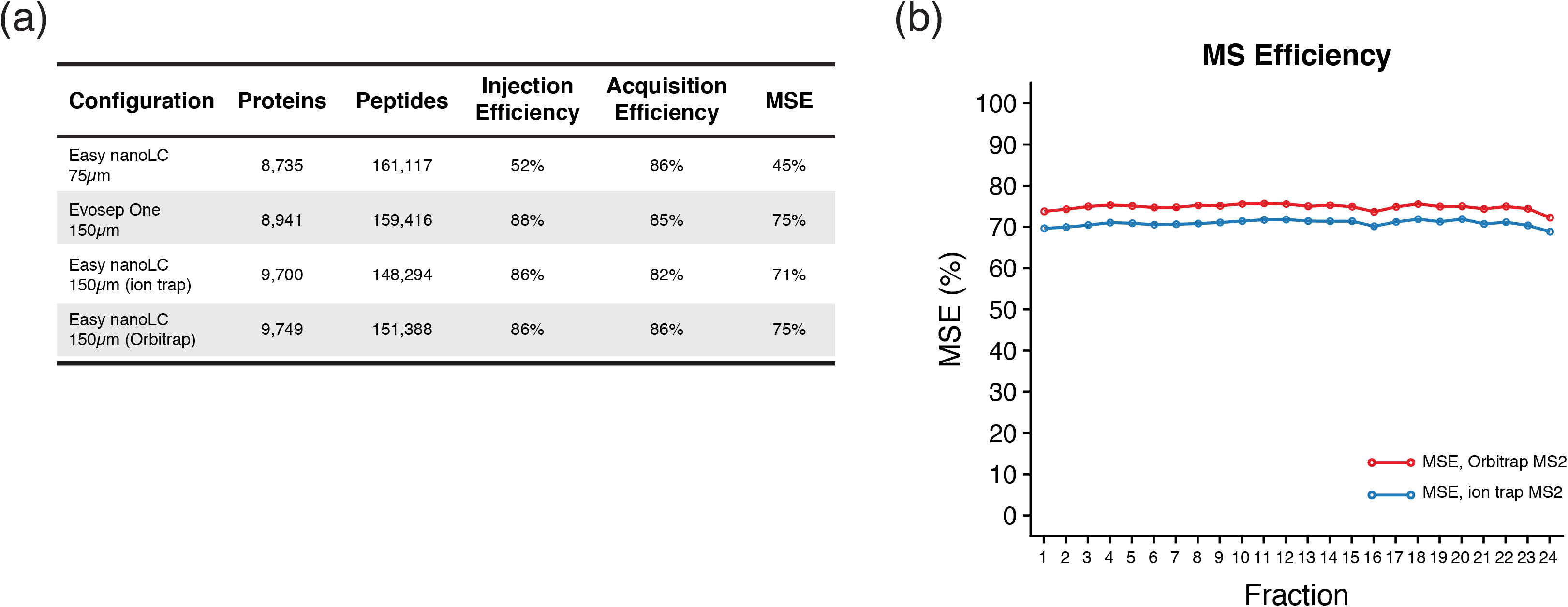
In-depth proteome analysis with a high MS efficiency is capable on a standard Easy nanoLC system. Peptides derived from an HEK293 lysate were subjected to offline fractionation with concatenation prior to analysis on an Easy nanoLC system equipped with a 150 μm ID column. Each of the final 24 fractions were analyzed for a total of 60-minutes each (including LC overhead). Data were acquired on an Orbitrap Fusion Lumos using either ion trap or Orbitrap MS2 detection. **(a)** Table displaying protein and peptide identification metrics along with calculated efficiency values. Data acquired in this study are displayed alongside values from an Easy nanoLC (75 μm ID column) and an Evosep One obtained from a previously published analysis. **(b)** Line plot displaying MSE values calculated across all fractions from the ion trap and Orbitrap data acquired in this study.

In this work, the ability to improve MS utilization performance when utilizing a standard nano-LC system is demonstrated. Equipping an Easy nanoLC system with a larger 150 μm ID column resulted in substantially reduced injection overheads. Although a reduction in sensitivity was observed in comparison to smaller 50 μm ID columns, the signal loss was mitigated by leveraging the increased injection capacity of the larger 150 μm column. The 150 μm column setup was then applied to the in-depth analysis of a mammalian cell proteome where a substantial improvement in MS efficiency was observed compared with previous results while illustrating no sacrifice in the obtained proteome coverage. Together, these data demonstrate that balancing LC overhead with fractionation-concatenation by utilizing nano-chromatography columns with larger IDs can substantially improve data acquisition on standard nano-LC hardware.

The primary caveat of this approach is that it requires the use of additional material per injection to maintain sensitivity. However, proteomics experiments that aim to perform deep profiling of a sample typically load anywhere from 100 – 1,000 μg of peptide material for offline fractionation prior to MS analysis. On the low end of this scale, assuming lossless processing and concatenation into a final set of 24-fractions, a 100 μg sample will result in each concatenated sample containing 4 μg of available peptide for injection. In our experience, a typical 150 μm ID nano-chromatography column will have a loading capacity of approximately 5-6 μg of peptide material before band broadening begins to be observed. Based on the above data, 2 μg of on-column material is sufficient to recover signal lost due to the use of a 150 μm ID column (versus 50 μm). Therefore, this setup is applicable in a wide range of proteomics analyses and is portable to virtually any nano-LC system.

## Supporting Information

The following files are available free of charge at ACS website http://pubs.acs.org:

- Supporting information describing the drop in MS2 quality and matching with increasing column ID (Figure S1). Supporting information relating to the visualization of empty elution windows in non-concatenated peptide fractions (Figure S2).

## Acknowledgements

C.S.H. would like to acknowledge valuable input and discussions from Lida Radan.

## Author Contributions

C.S.H. conceived the idea, carried out the data analysis, and wrote the manuscript. H.A. contributed to the experimental design. H.A., S.L., C.C., P.H.S., and G.B.M. contributed to writing of the manuscript.

