## Supplemental Figures for "High-flow Nano-chromatography Columns Facilitate Rapid and High-quality Proteome Analysis on Standard Nano-LC Hardware"

### Supporting Figures

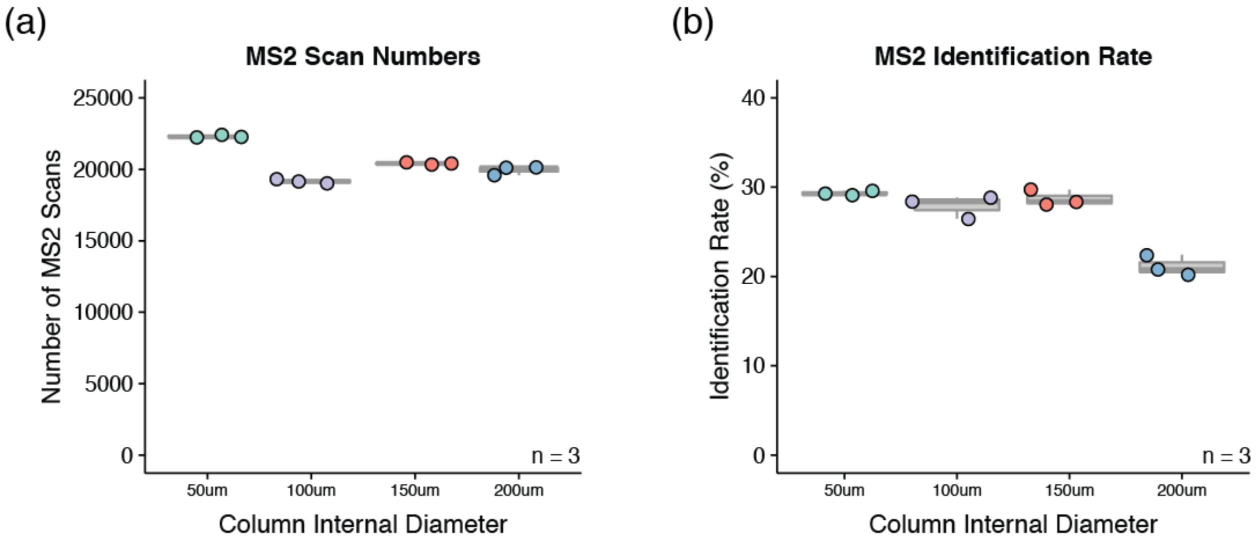

**Figure S1 – Nano-chromatography columns with larger IDs reduce the acquisition and identification rate from MS2 spectra.** Peptides derived from an HEK293 lysate were subjected to injection replicate (n = 3) analyses on an Easy nanoLC system equipped with columns of increasing ID (50 µm, 100 µm, 150 µm, 200 µm). Each sample was analyzed by an Orbitrap Fusion Lumos for 30 minutes using the same gradient conditions. **(a)** Boxplot depicting the number of MS2 spectra acquired in each of the replicate injections. **(b)** Boxplot depicting the peptide identification rate from MS2 spectra acquired in each analysis. Identification values are based on RawTools QC analysis using X!Tandem.

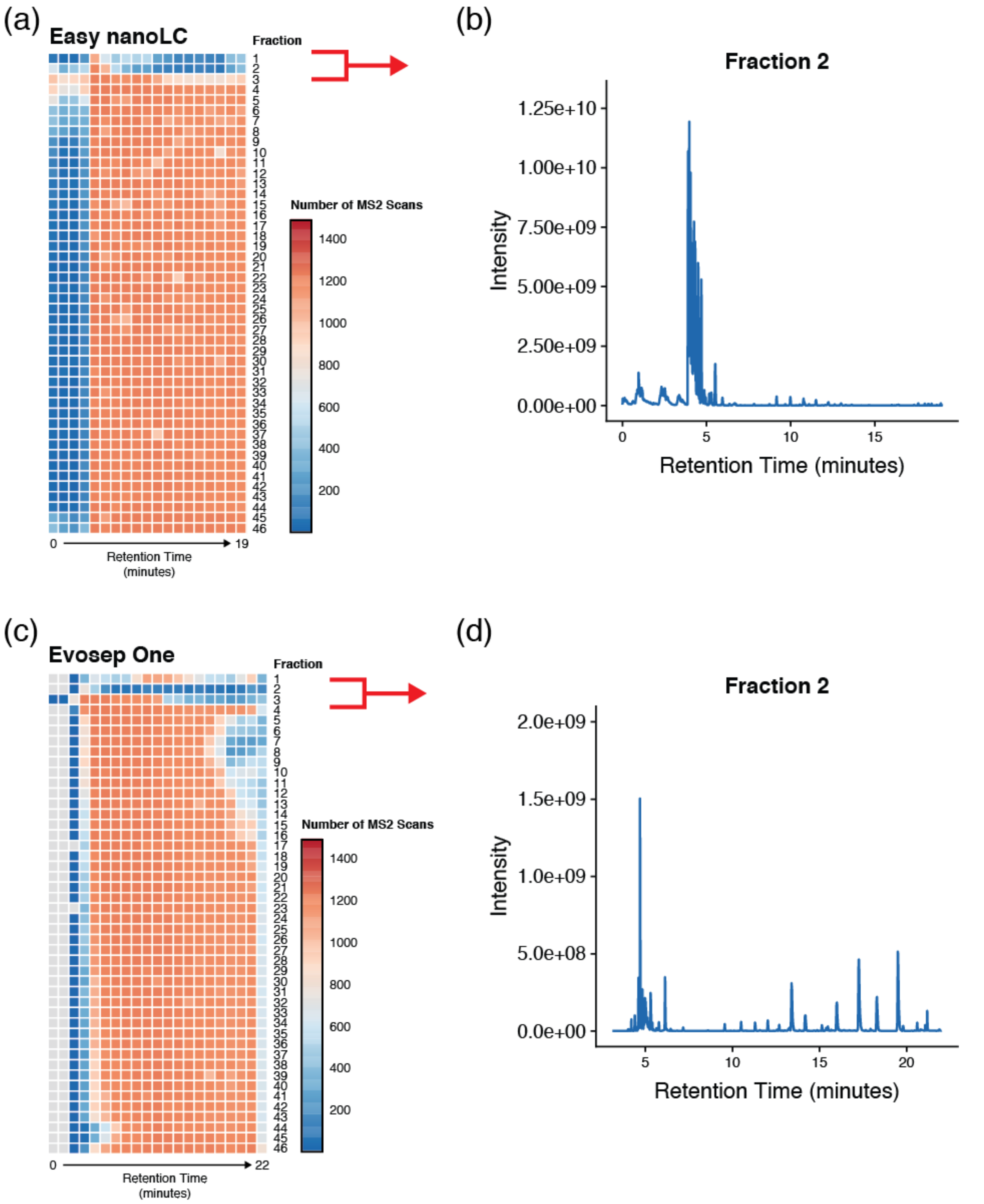

**Figure S2 – Non-concatenated fractions display low density elution windows.** Previously published data from HeLa cell peptides was acquired using a Q-Exactive HF-X system equipped with an Easy nanoLC or Evosep One nano-LC. Samples were separated into 46 fractions and analyzed without concatenation. Fraction data were analyzed with RawTools to generate metrics and chromatogram data. **(a)** Heatmap of the number of MS2 scans triggered in 1-minute bins across the entire elution window for each of the analyzed fractions from the Easy nanoLC data. **(b)** Base peak chromatogram of the second fraction from the analysis with the Easy nanoLC system. **(c)** Heatmap of the number of MS2 scans triggered in 1-minute bins across the entire elution window for each of the analyzed fractions from the Easy nanoLC data. **(d)** Base peak chromatogram of the second fraction from the analysis with the Evosep One system.
